# Spindle Oscillations Emerge at the Critical State of Electrically Coupled Networks in the Thalamic Reticular Nucleus

**DOI:** 10.1101/2023.12.31.573769

**Authors:** Shangyang Li, Chaoming Wang, Si Wu

**Affiliations:** School of Psychological and Cognitive Sciences, Beijing Key Laboratory of Behavior and Mental Health,, IDG/McGovern Institute for Brain Research, Center of Quantitative Biology,, Peking-Tsinghua Center for Life Sciences, Academy for Advanced Interdisciplinary Studies, Peking University, Beijing, 100871, China; Guangdong Institute of Intelligence Science and Technology, Hengqin, Zhuhai, 519031, Guangdong, China

**Keywords:** Spindle Oscillation, Electrical Synapse, Neuronal Heterogeneity, Critical State, Thalamic Reticular Nucleus

## Abstract

Spindle oscillation is a waxing-and-waning neural oscillation observed in the brain, initiated at the thalamic reticular nucleus (TRN) and typically occurring at 7-15 Hz. Experiments have shown that in the adult brain, electrical synapses, rather than chemical synapses, dominate between TRN neurons, suggesting that the traditional view of spindle generation via chemical synapses may need reconsideration. Based on known experimental data, we develop a computational model of the TRN network, where heterogeneous neurons are connected by electrical synapses. The model shows that the interplay between synchronizing electrical synapses and desynchronizing heterogeneity leads to multiple synchronized clusters with slightly different oscillation frequencies, whose summed activity produces spindle oscillation as seen in local field potentials. Our results suggest that during spindle oscillation, the network operates at the critical state, which is known for facilitating efficient information processing. This study provides insights into the underlying mechanism of spindle oscillation and its functional significance.

## INTRODUCTION

Spindle oscillation refers to a waxing-and-waning oscillatory event occurring at a frequency of 7-15 Hz^1,2^. This phenomenon has been widely observed in various mammals, including the cat^3,4^, mice^5,6^, rat^7,8^, ferret^9,10^, dog^11,12^, sheep^13^, macaque^14,15^, and human^16,17^. Extensive experimental studies have established a strong correlation between spindle oscillation and learning and memory processes in the brain^18^. For example, research has shown that the density and amplitude of spindle oscillation increase during post-learning sleep, both for declarative memory tasks^19–21^ and procedural memory tasks^22,23^. Manipulation studies conducted on humans and rodents have provided causal evidence supporting the role of spindle oscillation in memory consolidation^24,25^. Despite the recognized importance of spindle oscillation in brain functions, the neural mechanism underlying spindle generation remains a subject of debate.

The conventional understanding of spindle generation has suggested that spindles are generated in the thalamus^1,2,17,26^. For example, early investigations on anesthetized cats revealed that the decorticated thalamus alone could produce waxing-and-waning spindle rhythms^27,28^. In vitro studies using thalamic slices further demonstrated that the synaptic connectivity between thalamocortical (TC) relay neurons and thalamic reticular nucleus (TRN) neurons played a crucial role in spindle rhythm generation^9,10,29–32^. According to this framework, low-threshold bursts from TRN cells transiently inhibit TC neurons, and upon release from inhibition, TC cells generate Ca^2+^-dependent post-inhibitory rebound bursts, which drive subsequent spike bursts in TRN neurons^26,33–36^. However, experimental findings have challenged this “ping-pong” model of spindle generation. These inconsistencies include: 1) optogenetic stimulation of TRN neurons triggering cortical spindles that are not observed in the corresponding TC site^37^; 2) the generation of sleep spindles in the mouse thalamus in the absence of T-type Ca^2+^ channels and burst firing of TC neurons, both in vivo and in vitro^38^; and 3) administration of a Ca^2+^ channel blocker in the corticothalamic network reducing spindles at higher frequencies (approximately 14 Hz) while enhancing the power of slow spindles (approximately 10 Hz)^39^. These findings strongly suggest that the involvement of TC cells is not the necessary condition for spindle generation^33^.

TRN neurons are intrinsic GABAergic cells that encompass the entire thalamus^40,41^. They receive glutamate excitatory inputs from TC and corticothalamic (CT) neurons^40,42^, as well as extensive inhibitory projections from subcortical areas, including the basal forebrain^43–46^, the substantia nigra pars reticulata^47^, the globus pallidus^48–50^, and the hypothalamus^51^. Experimental studies have suggested that the TRN can generate oscillations at the spindle frequency without relying on the inputs from the cortex and thalamus^4,52,53^. Earlier experiments involving local glutamate application revealed that TRN neurons in rodents younger than 2 weeks of age are interconnected through inhibitory chemical synapses^54–58^. Subsequent computational modeling demonstrated that a TRN network with neurons connected by inhibitory chemical synapses can generate various types of network oscillations at frequencies of 7-10 Hz^34,59–61^. However, these models rely on low-threshold Ca^2+^-dependent bursts, which either require a sufficiently negative reversal potential of inhibitory synapses^34,59,60^ or a hyperpolarized resting membrane potential^61^. These assumptions are not consistent with the biological properties of TRN neurons, which include: 1) TRN cells have a depolarized chloride reversal potential (E_GABA_ = − 62 mV) compared to TC neurons, resulting in postsynaptic depolarizations rather than strong inhibition upon receiving GABAergic inputs^62–64^. 2) Spindles can be generated from TRN slices with a more depolarized resting potential (−63.8 ± 1.2 mV)^29,31^.

Moreover, experimental findings have shown that electrical synapses dominate the connectivity between TRN neurons after two weeks of age. Paired recordings from TRN neurons in rats and mice across different ages have revealed the presence of electrical synapses but rare chemical transmission between TRN neurons^65–69^. Optogenetic stimulation of layer 6 CT neurons or neighboring TRN neurons failed to evoke inhibitory postsynaptic currents (IPSCs) in TRN neurons in mice older than two weeks^70–72^. Genetic deletion of vesicular GABA transporters from TRN neurons only reduced GABAergic transmission in TC neurons but not TRN neurons^72^. These findings indicate that the electrically coupled TRN network, without direct inputs from TC neurons, is sufficient for the generation of spindle oscillations. Notably, sleep spindles begin to emerge around 2-3 months of age in humans and during the third to fourth postnatal weeks in rodents^73^, which aligns with the dominance of electrical couplings in the TRN at later developmental stages. Thus, the traditional mechanism based on chemical synapses for spindle generation in the TRN suggests a reconsideration.

Motivated by recent experimental evidence, the present study investigates the generation of spindle oscillation by the electrically coupled TRN neurons. Here, sleep spindle oscillations in the TRN are defined as oscillatory events in the frequency range of 7-15 Hz and with a typical duration of 0.5-3 seconds. They are characterized by a waxing-and-waning pattern in the os-cillatory activity, as identified through visual inspection of the waveform filtered between 0.3-35 Hz. We develop a computational model that incorporates the latest findings regarding TRN neurons, specifically the prevalence of electrical synapses between neurons and the presence of neuronal heterogeneity. These factors have opposing effects on network dynamics, as electrical coupling tends to synchronize neurons while neuronal heterogeneity tends to desynchronize them. Our results suggest that the interplay between electrical coupling and neuronal heterogeneity leads to the self-organization of the TRN network into the critical state. In this state, neurons form multiple synchronized clusters, each oscillating at slightly different frequencies. The superposition of these neuronal activities, mimicking the local field potential (LFP) recorded in experiments, exhibits characteristic spindle oscillations. Overall, our modeling study suggests a natural emergence of spindle oscillations in the TRN network when it operates in the critical state — a condition known to facilitate efficient information processing in complex systems^74^. These findings provide insights into the underlying mechanisms of spindle generation and offer further understanding of the functional role of the TRN in regulating thalamocortical activity.

## RESULTS

### An electrically coupled heterogeneous TRN network

We built a TRN network model based on physiological data collected from ferrets, cats, mice, and rats. Specifically, we considered that the TRN network is a two-dimensional grid (Figure 1A), on which *N* neurons are uniformly distributed and connected with each other by electrical synapses^65,66,68,69,72^ according to a Gaussian probability connection profile (this reflects the fact that neighboring TRN neurons tend to be connected by electrical synapses^68^, see details in STAR). Each neuron in the network receives two kinds of external synaptic inputs: excitatory synaptic projections from the thalamus and cortex^40^, and inhibitory inputs from subcortical brain areas^43–51^. The dynamics of a single TRN neuron is given by^75^,

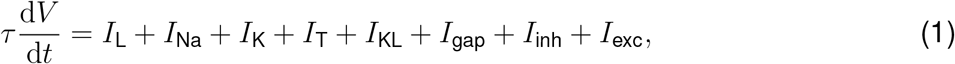

where *V* is the membrane potential and *τ* the membrane time constant. *I*_*L*_, *I*_Na_, *I*_K_, *I*_T_ and *I*_KL_ denote, respectively, the leaky channel current, the sodium channel current, the potassium channel current, the low-threshold calcium channel current, and the potassium leaky current. *I*_gap_ denotes the recurrent gap junctional current from other TRN neurons. *I*_inh_ and *I*_exc_ denote the subcortical inhibitory current^50,51^ and the thalamic and cortical excitatory current^40,42^, respectively. For the detailed form of each current, see STAR.

**Figure 1.**
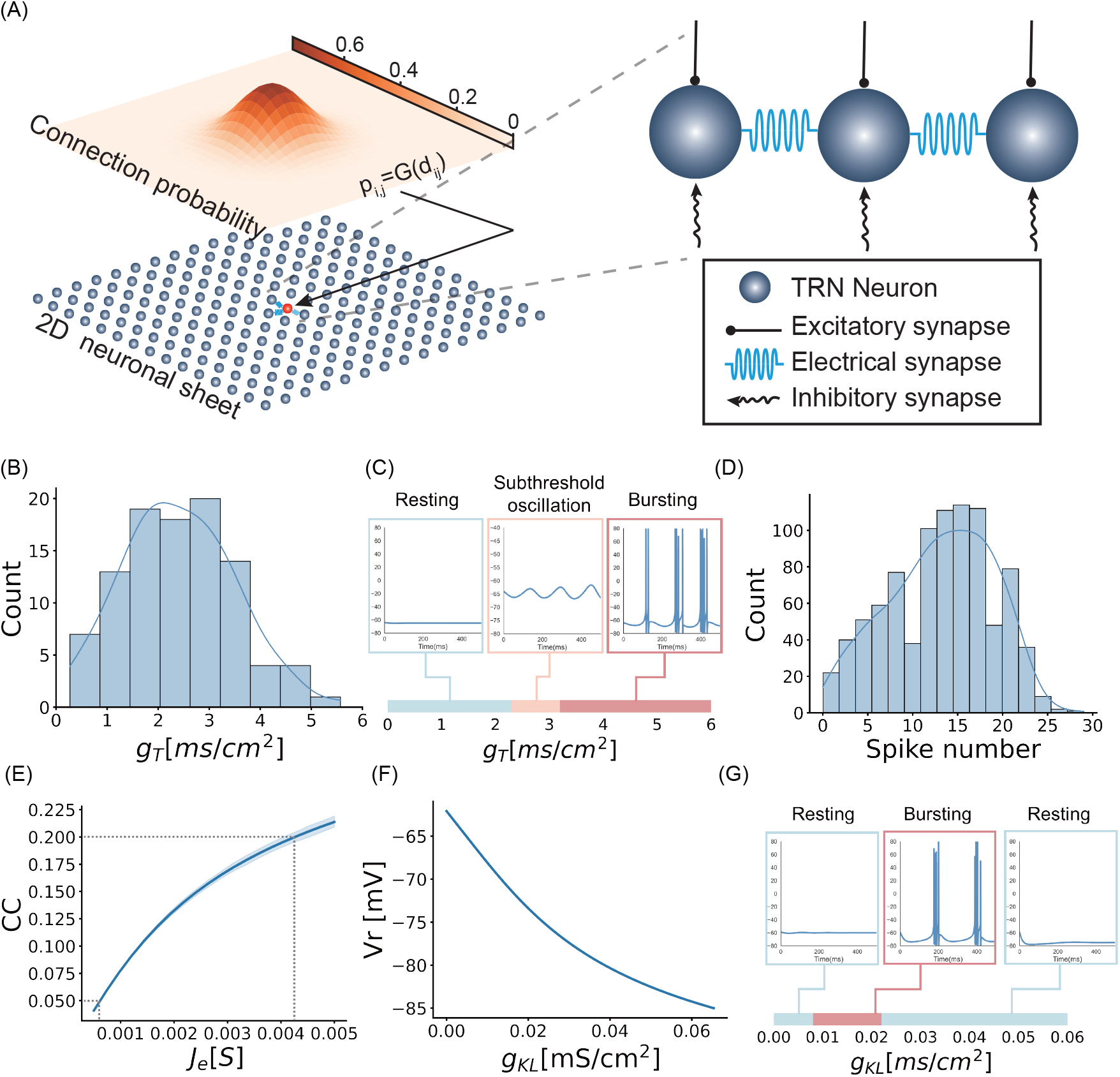
The TRN network model. (A) The network structure. TRN neurons are uniformly distributed on a two-dimensional sheet and connected with electrical synapses. The connection probability between neurons follows a distance-dependent Gaussian distribution. Each neuron receives an excitatory thalamic and cortical input and an inhibitory subcortical input.. (B) The distribution of the conductance *g*_*T*_ of T-type Ca^2+^ channels of all TRN neurons.. (C) The spontaneous activity of a TRN neuron changes from the resting state, subthreshold oscillation, to bursting firing when increasing the conductance *g*_*T*_.. (D) The distribution of the number of spikes per burst in TRN neurons. Each burst was triggered by injecting a negative current followed by abrupt current removal^83^.. (E) The relationship between the coupling coefficient (CC) and the conductance of electrical synapse *J*_*e*_. For CC in the range of (0.05 − 0.2), *J*_*e*_ is in the range of (0.0005*S*, 0.005*S*). The shaded region represents the standard error of the mean (SEM) calculated from *n* = 10 independent simulations.. (F) The relationship between the resting membrane potential and the potassium leaky conductance *g*_*KL*_.. (G) The spontaneous activity of a TRN neuron changes from the resting state, bursting firing, to the resting state when increasing the neuromodulation level *g*_*KL*_.

In the experiments, spindle oscillation was observed in the local field potential (LFP). Similar to previous works^76^, we considered that the simulated LFP in our model represents the averaged membrane potential of all TRN neurons, which is calculated to be,

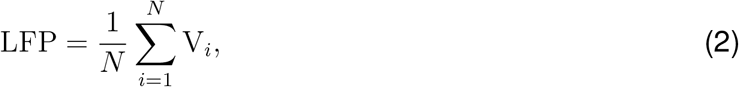

where *V*_*i*_ is the membrane potential of the *i*-th neuron, and the summation runs over all TRN neurons. We also calculated LFP using a different method called the spatial lowpass filter and obtained the same results (see STAR).

In our model, the generation of spindle oscillation is influenced by three crucial factors: neuron heterogeneity, electrical couplings between neurons, and neuromodulation levels. In the below, we will discuss the settings of these factors in the model and explore their effects on the neural dynamics.

### Heterogeneity of TRN neurons

TRN neurons are distinguished by their abundant T-type Ca^2+^ channels^77^. In our model, we computed the T-type Ca^2+^ channel current using the equation *I*_*T*_ = *g*_*T*_ *f* (*V*) (refer to Equation 5 for the specific form of *f* (*V*)), where *g*_*T*_ represents the channel conductance. It is important to note that TRN neurons exhibit significant heterogeneity. Different TRN neurons display distinctive firing patterns, which have been linked to variations in sleep rhythms across sensory cortices^78,79^. One of the primary sources of this heterogeneity is the varied distribution of T-type Ca^2+^ channels among TRN neurons^78–83^. Therefore, in our model, we assigned different values of *g*_*T*_ to individual neurons to reflect this heterogeneity (see Figure 1B).

We explored how *g*_*T*_ affects the dynamics of a single neuron. By varying *g*_*T*_, we observed that (Figure 1C): 1) for *g*_*T*_ *<* 2.32 mS*/*cm^2^, the neuron is in the resting state; 2) for 2.32 mS*/*cm^2^ *< g*_*T*_ *<* 3.19 mS*/*cm^2^, subthreshold oscillation occurs as seen in the voltage clamp experiment^66^, and the amplitude of subthreshold oscillation increases with *g*_*T*_ ; 3) for *g*_*T*_ *>* 3.19 mS*/*cm^2^, the neuron generates bursting spikes.

To match with the experimental data, we considered that the values of *g*_*T*_ for all neurons are randomly sampled from a truncated normal distribution as illustrated in Figure 1B. In the biological experiment, the number of spikes per offset burst of TRN neurons was measured and was found to approximately satisfy a normal distribution (Fig.3d in^83^). Following the same protocol as in^83^, we injected a negative current to a TRN neuron followed by a sudden current removal to induce bursting firing of the neuron. We set the distribution of *g*_*T*_ appropriately, such that the numbers of spikes in bursts generated by all TRN neurons exhibit a similar distribution as observed in the experiment (Figure 1D).

### Electrical couplings between TRN neurons

Another remarkable feature of TRN neurons is their electrical couplings^65,66,72^. In the model, we set the electrical synapse current between neurons (*i, j*) to be *I*_*gap*_ = *J*_*e*_(*V*_*j*_ − *V*_*i*_) (see STAR for details), with *J*_*e*_ the coupling conductance. The strength of electrical couplings can be quantified by the coupling coefficient (CC) between TRN neurons. To correlate the electrical synapse conductance *J*_*e*_ with the CC, we calculated how the average CC varies with *J*_*e*_ (see Figure 1E). To be consistent with the experimental finding that the average CC in TRN varies in the range of (0.05 − 0.2)^66,84^, we set the conductance *J*_*e*_ of electrical synapse in the range of (0.0005 *S*, 0.005 *S*) in all later computational experiments.

### Modulation by neuromodulators

It is known that subcortical neuromodulators, such as acetylcholine and norepinephrine, influence oscillatory activities in the thalamus^85,86^. To model this neuromodulation effect, we included a potassium leaky current *I*_*KL*_ = *g*_*KL*_(*E*_*KL*_ − *V*) in the neuronal dynamics (see Equation 5 for more details), where *g*_*KL*_ is the conductance controlled by neuromodulators. At the single neuron level, *g*_KL_ shows a great impact on the membrane resting potential, i.e., the value of resting potential decreases with *g*_KL_ (Figure 1G). Neuromodulation also affects neuronal firing patterns. As illustrated in Figure 1F, a TRN neuron tends to oscillate at an intermediate *g*_KL_ value and rest when *g*_KL_ is small or large.

#### Spindle oscillation emerges in the TRN network

Experimental studies have revealed varied oscillation behaviors in the TRN network during different sleep and awake conditions^87^. To validate our network model, we first identify the parameter ranges in which TRN spindles occur. Specifically, we manipulate the parameter *g*_KL_, which reflects the levels of neuromodulation in different sleep and awake conditions, while keeping all other parameters constant at appropriate values (see STAR). Furthermore, when modeling the sleep condition, we incorporate an inhibitory current *I*_inh_ from subcortical regions, while excluding any excitatory current *I*_exc_ from the thalamic and cortical regions. Conversely, in the awake condition, we include both *I*_inh_ and *I*_exc_.

### Deep sleep condition

In deep sleep, i.e., N3 NREM sleep, the neuromodulation level is low^88^, we set *g*_KL_ to be a large value. In such a state, most TRN neurons are hyperpolarized and they predominantly generate strong bursting activities with high-frequency inter-burst spikes (Figure 2A). Together with electrical couplings between neurons, the whole network generates highly synchronized responses, resulting in strong LFP (Figure 2D), and the oscillation frequency is around 4 Hz (Figure 2G), agreeing with the delta oscillation (1 − 4 Hz) observed in the experiment^89^.

**Figure 2.**
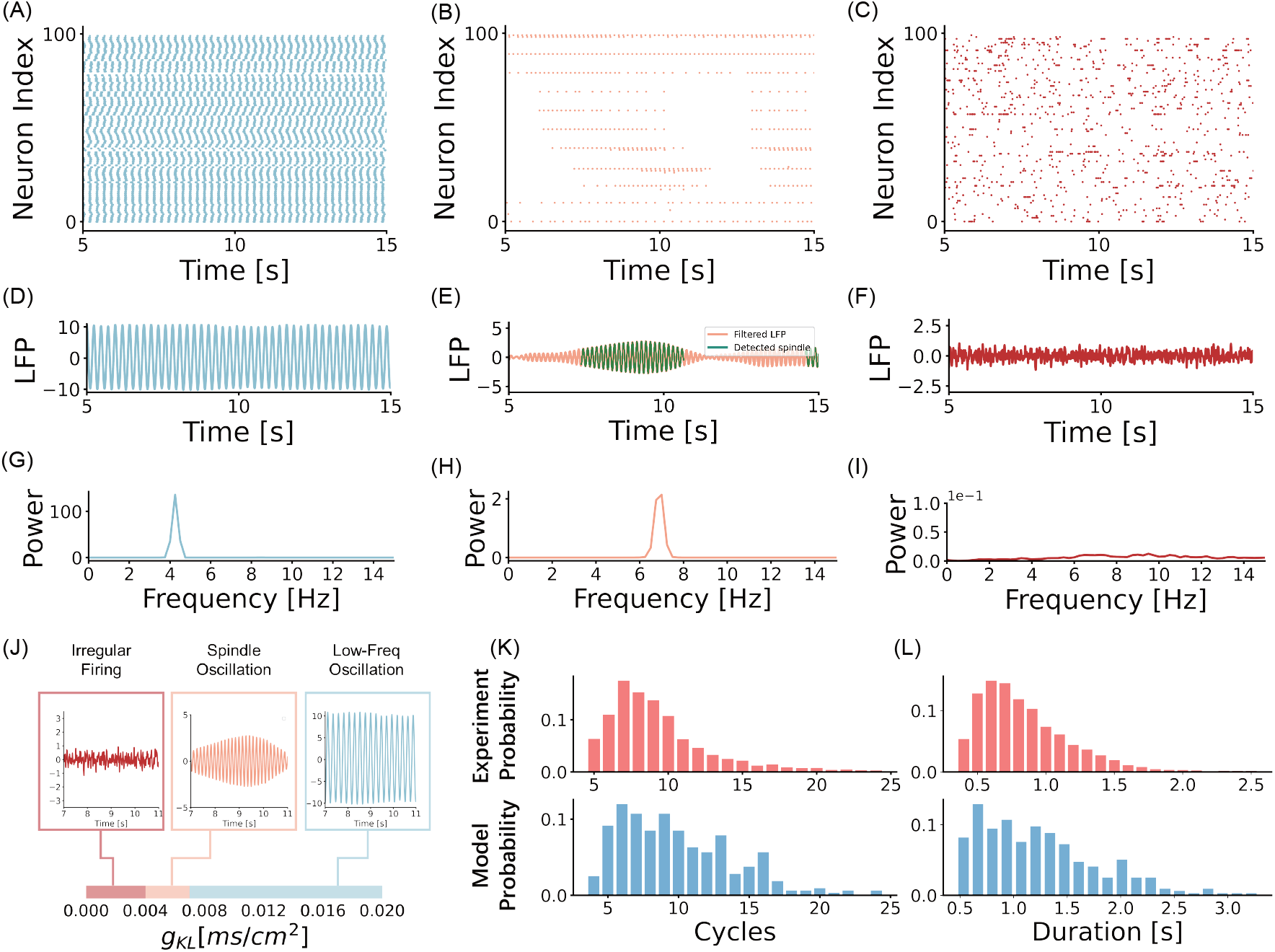
Oscillation behaviors of the TRN Network. (A-C) Raster plots of neurons’ firings in deep sleep, light sleep, and alertness conditions, respectively.. (D-F) LFP in deep sleep, light sleep, and alertness conditions, respectively.. (G-I) The spectrum of LFP in deep sleep, light sleep, and alertness conditions, respectively.. (J) The phase diagram of the network state with respect to the neuromodulation level *g*_*KL*_.. (K) The statistical properties of the spindle oscillation cycle in experiments^7^ and our model. Upper panel: the experimental data. Lower panel: the model simulation data.. (L) The statistical properties of the spindle oscillation duration in experiments^7^ and our model. Upper panel: the experimental data. Lower panel: the model simulation data.

### Light sleep condition: spindle oscillation occurs

In light sleep, i.e., N2 NREM sleep, the neuromodulation level is at the medium level^88^, we set *g*_KL_ to be a mediate value. In this state, TRN neurons are less hyperpolarized (−65.5 ∼ −63.5 mV) and produce weak bursts (Figure 2B). A single burst typically contains only one or two spikes, and such bursts can be easily interrupted (see the discontinuous spiking of a TRN neuron in Figure 2B). Under this condition, the whole TRN network generates oscillations of the waxing-and-waning shape in LFP (Figure 2E) with a frequency of 7 Hz (Figure 2H), which is in the range of spindle oscillation (7 − 15 Hz)^18^. We also calculated LFP using a different method and obtained the similar results (see Figure S12).

We carried out ablation studies to illustrate the impact of model parameters on spindle generation. First, by removing the subcortical synaptic inputs (i.e, setting *I*_inh_ = 0), the waxing-and-waning oscillation disappears in the network (see Figure S10). Second, by removing or blocking the low-threshold Ca^2+^ channel currents (i.e., setting *I*_*T*_ = 0), spindle oscillation also disappears (see Figure S11). These results are in line with the experimental findings, which are: 1) the TRN slice *in vitro* without subcortical projections could not produce spindle oscillation^31^; 2) the power of spindle oscillations was significantly reduced in Ca^2+^ channel knockout mice, and the spindle rhythmogenesis in major sensory cortical areas depended on low-threshold Ca^2+^ currents^79,90,91^.

### Alertness condition

In the alertness condition, i.e., REM sleep or awake state, the neuromodulation level is high^88^, we set *g*_KL_ to be small. In such a case, TRN neurons have depolarized resting membrane potentials (Figure 1C). Furthermore, in the REM sleep state, we set the excitatory input *I*_exc_ = 0. Consistent with experimental finding^88^, a significantly weak LFP signal is seen in our TRN network (Figure S4B). Moreover, TRN neurons show small voltage fluctuations and uncorrelated activities (Figure S4C) and TRN neurons under such a state present nearly no spiking activity. In the awake state, we set *I*_exc_ ≠ 0 representing the excitatory input from the thalamus and cortex, and we found that TRN neurons show irregular spiking (Figure 2F) and have no dominating oscillation frequency (Figure 2I).

### Phase diagram of the TRN network

In summary, Figure 2J presents how the TRN network undergoes three different states of oscillation as a function of the neuromodulation level parameter *g*_KL_. As *g*_KL_ decreases (i.e., the neuromodulation level increases), the TRN network transitions from the regular low-frequency delta oscillation (3 −6 Hz), to the waxing-and-waning spindle oscillation (6 − 8 Hz), and finally to irregular firings. These states correspond to the physiological states of deep sleep, light sleep, and alertness, respectively. The resting membrane potentials of individual neurons also change accordingly, rising from (−65.5 ∼ −75mV), to (−65.5 ∼ −63.5mV), and to (*>* −63.5 mV). Our model reproduces the experimental finding^52^ that the resting membrane potential in spindle oscillation is more depolarized than that in delta oscillation.

### Statistical properties of spindle oscillations

The experimental data has shown that spindle oscillations are characterized as 0.5 − 3s long oscillation sequences in LFP which have the waxing-and-waning profile and oscillate at 7 − 15 Hz^18^. The density, duration, and amplitude of spindle oscillations, all display statistical characteristics, and they are highly correlated with memory performance^92^, problem-solving ability^93^, and mental diseases^94^. We found that our model can reproduce the statistical properties of spindle oscillations observed in human^95–97^ and animal^7,38^. By using multichannel silicon-probe recording on the thalamus of urethane-anesthetized and sleeping rats, it was found that both the spindle oscillation cycle and the duration exhibit uni-modal distributions with biases to small values (refer to the upper panels in Figure 2K and L)^7^. Consistent with these results, our network model generates spindle oscillations having similar statistical distributions (refer to the lower panels in Figure 2K and L), with the majority of spindle oscillations having 5 −13 oscillation cycles and a duration of 0.5 − 1.5s. These results are also in line with the observations in human^95^ and mice^38^.

#### The mechanism of spindle generation

In the below, we further explore the underlying mechanism of spindle generation in our TRN network. Specifically, we show that the interplay between electrical coupling and neuronal heterogeneity produces spindle oscillations.

#### Effects of electrical coupling

The role of electrical coupling in regulating TRN oscillation has been highlighted in the experiments^98,99^. Here, we systematically investigated the effects of electrical coupling on the dynamics of the TRN network. We fixed all other parameters of the model except the conductance of electrical synapse *J*_*e*_. The parameter *g*_KL_ reflecting neuromodulation is set to be of an intermediate value which allows the generation of spindle oscillation (see Section).

We first examined the relationship between the conductance of electrical synapse *J*_*e*_ and the cross-correlation (CC) among neurons in the network. As depicted in Figure S6A, we observed that CC increases as *J*_*e*_ rises. This phenomenon can be attributed to the synchronization of neuronal responses facilitated by electrical couplings. To further explore the impact of *J*_*e*_ on spindle generation, we conducted measurements and documented the probability of spindle occurrence at different *J*_*e*_ values (refer to STAR). The results, illustrated in Figure S6B, reveal an initial increase followed by a subsequent decrease in the probability of spindle generation with increasing *J*_*e*_. This pattern suggests that an appropriate range of electrical coupling strength is required for the spindle generation. This observation aligns with our intuition: when *J*_*e*_ is too small, neurons in the network fire independently, resulting in a lack of coherent oscillation in the local field potential (LFP); conversely, when *J*_*e*_ is too large, all neurons fire synchronously, leading to LFP oscillation at a single frequency. The subsequent sections provide a more detailed analysis of the mechanics underlying spindle generation.

We further conducted an analysis on the clustering phenomenon within the network. In this context, a cluster represents a group of interconnected neurons that fire at the same frequency. Our observations reveal that by increasing the electrical coupling conductance (*J*_*e*_), the network state can be categorized into three distinct phases (Figure 3). These phases include: 1) the asynchronous firing phase (AFP, depicted in Figure 3A-C), where neuronal firings are largely independent of each other, the network lacks obvious large firing clusters, and the LFP does not exhibit clear oscillation during this phase; 2) the multiple cluster phase (MCP, illustrated in Figure 3D-F), characterized by the formation of multiple clusters within the network, varying in size and oscillating at slightly different frequencies. The LFP during this phase displays spindle oscillation; 3) the synchronized firing phase (SFP, shown in Figure 3G-I), where the majority of neurons in the network synchronize their firing, resulting in the LFP oscillating at a single dominant frequency. Notably, the occurrence of spindle oscillation is associated with the MCP, and this phenomenon of multiple clustering has also been reported in a current clamp experiment conducted on the rat TRN slice^66^.

**Figure 3.**
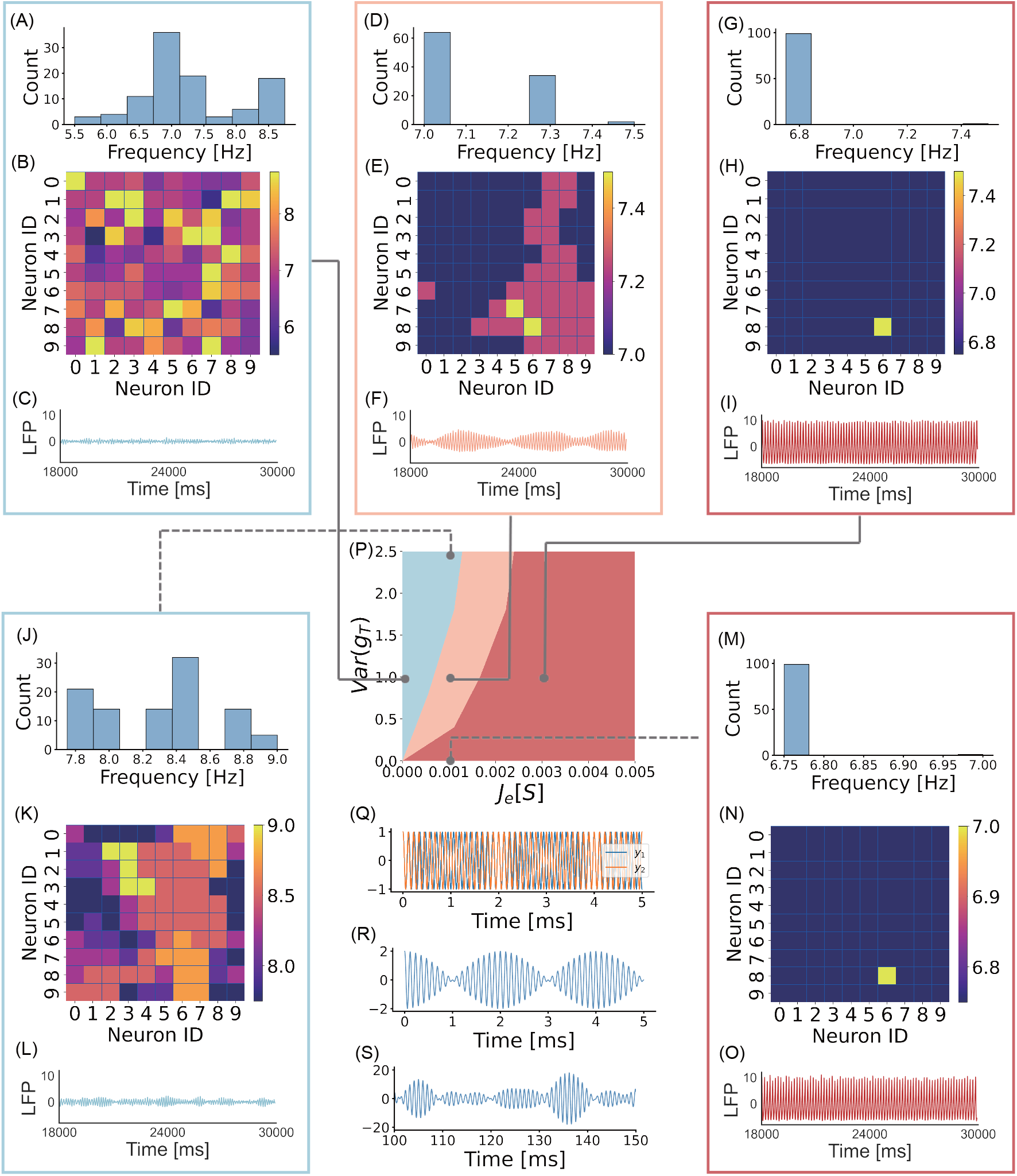
The mechanism of spindle generation. (A-C) Histogram of neuronal oscillation frequency, heat map of neuronal oscillation frequency, and LFP signal. The parameters are: *J*_*e*_ = 0 and Var(*g*_*T*_) = 1.0.. (D-E) The same as in (A-C), except *J*_*e*_ = 0.001 and Var(*g*_*T*_) = 1.0.. (G-I) The same as in (A-C), except *J*_*e*_ = 0.003 and Var(*g*_*T*_) = 1.0.. (J-L) The same as in (A-C), except *J*_*e*_ = 0.001 and Var(*g*_*T*_) = 2.4.. (M-O) The same as in (A-C), except *J*_*e*_ = 0.001 and Var(*g*_*T*_) = 0.. (P) Phase diagram of the TRN network with respect to the electrical coupling strength *J*_*e*_ and the neuronal heterogeneity Var(*g*_*T*_).. (Q) Two sine waves having slightly different oscillation frequencies, where *y*_1_(*t*) = sin(2.2*πt*) and *y*_2_(*t*) = sin(2.4*πt*).. (R) Superposition of two sine waves, *y*(*t*) = *y*_1_(*t*) + *y*_2_(*t*).. (S) Superposition of 100 sine waves by their frequencies corresponding to the oscillation fre-quencies of TRN neurons during spindle generation.

In the above, we have assumed a constant electrical synapse conductance between all neurons for simplicity. However, empirical evidence suggests that the strength of electrical synapses between TRN neurons exhibit significant variability^100,101^. To investigate the impact of electrical coupling variability on spindle oscillations, we conducted an additional experiment incorporating heterogeneous synapse strengths. We modeled the electrical synapse strengths using an exponential decay distribution, reflecting the biological tendency for spatially proximate neurons to form stronger connections (see Figure S5A and B). Despite this added heterogeneity, our model continued to generate robust spindle oscillations (see Figure S5C and E). This finding highlights that our simplified models effectively capture the essential role of electrical synapses in spindle generation.

#### Effects of neuronal heterogeneity

We conducted further investigation into the impact of neuronal heterogeneity on the dynamics of the TRN network. Neuronal heterogeneity, quantified by the variance of the T-type Ca^2+^ conductance (*g*_*T*_) denoted as Var(*g*_*T*_) (while keeping the mean of *g*_*T*_ fixed), plays a significant role in shaping the distinctive behavior of individual neurons and contributes to the desynchronization of their activities. This is evident in the decrease of cross-correlation between neurons as Var(*g*_*T*_) increases (Figure S6C). We also observed that the probability of spindle generation initially increases with Var(*g*_*T*_) and then decreases (Figure S6D). This observation can be intuitively understood. When Var(*g*_*T*_) is very small, neurons in the network exhibit synchronous firing, resulting in a single-frequency oscillation in the LFP. Conversely, when Var(*g*_*T*_) is very large, neurons fire independently, leading to the absence of coherent oscillation in the LFP.

Similar to the impact of electrical coupling, the clustering phenomenon within the TRN network also undergoes three distinct phases in response to neuronal heterogeneity. However, in the case of heterogeneity, these phases appear in the reverse order as Var(*g*_*T*_) increases, namely, SFP (Figure 3M-O), MCP (Figure 3D-F), and AFP (Figure 3J-L). The reason behind this ordering is straightforward: as the heterogeneity represented by Var(*g*_*T*_) increases, neuronal firings become progressively more desynchronized, resulting in a transition from a single cluster to multiple clusters, and eventually to the absence of clustering. Once again, we observed that spindle oscillations occur specifically during the MCP.

We also inspected the statistical properties of spindles when the neuronal heterogeneity varies. With the decrease of Var(*g*_*T*_), the averaged cluster size (i.e., the averaged number of neurons in a synchronized cluster) increases. Figure S7A displays how the spindle probability (measured as the number of spindles occurring in a fixed time interval) varies with the averaged cluster size. It shows that the spindle probability takes the maximum value when the averaged cluster size has an intermediate value, i.e., the neuronal heterogeneity Var(*g*_*T*_) takes an intermediate value. With the increase of Var(*g*_*T*_), the variance of individual neuronal oscillation frequency increases. Figure S7B shows that the spindle modulation frequency increases with the variance of neuronal oscillation frequency. This can be intuitively understood from the below simple mathematical analysis (see Equation 3), which shows that the spindle modulation frequency is determined by the difference between neuronal oscillation frequency. The above two statistical properties can be checked in experiments to validate our model.

Up to now, we have considered that the TRN neurons form a homogeneous group for simplicity. However, the experiments have identified three distinct tiers within the TRN, where central cells exhibit a higher tendency to burst, while edge cells favor tonic spiking^82,83,102,103^. To reflect this heterogeneity, we carried out an additional experiment, in which we extended the model by including distinct T-type calcium conductance profiles for edge and central cells. Specifically, the edge cells were represented by two 2×10 subnetworks, whereas the central cells were modeled as a 10×10 subnetwork (Figure S8A). We adjusted the means of *g*_KL_ in the edge and central tiers to align with the experimental measurements of the distribution of the number of spikes in rebound bursting (see Figure S8B and Figure. 3d in^83^). We confirmed that with this extra heterogeneity in the cell type distribution, our model can still produce robust spindle oscillations (Figure S8C-G), with spindles predominantly occurring in the central part of the network which has the higher mean value of *g*_KL_.

In summary, our results suggest that electrical coupling and neuronal heterogeneity exert contrasting effects on the dynamics of the TRN network. Electrical coupling promotes synchronization, whereas neuronal heterogeneity contributes to desynchronization. The interplay between these two factors gives rise to a diverse range of dynamics within the TRN network. Notably, when the amplitudes of electrical coupling and neuronal heterogeneity are appropriately balanced, spindle oscillation emerges. Figure 3P illustrates the phase diagram of the TRN network concerning the electrical coupling and neuronal heterogeneity.

#### Mathematical analysis of spindle generation

In the above, we have established that spindle oscillations emerge when the network state resides in the MCP. The MCP state is distinguished by the coexistence of multiple clusters with varying sizes and slightly different oscillation frequencies. In the below, we describe how the MCP state gives rise to the characteristic waxing-and-waning LFP pattern.

Let us first consider two harmonic oscillators with the same amplitude but slightly different frequencies (see the example illustrated in Figure 3Q), denoted, respectively, as *y*_1_(*t*) = *A* cos (2*πω*_1_*t*) and *y*_2_(*t*) = *A* cos (2*πω*_2_*t*), with |*ω*_1_ − *ω*_2_| ≪ (*ω*_1_ + *ω*_2_)*/*2. The superposition of the two oscillators *y*(*t*) = *y*_1_(*t*) + *y*_2_(*t*) can be expressed as

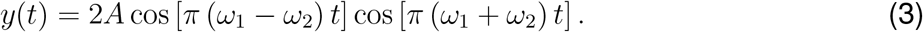

We see that the superposition of two oscillators displays a waxing-and-waning shaped oscillation. Specifically, the variable *y*(*t*) oscillates at a high frequency of (*ω*_1_ +*ω*_2_)*/*2, while its amplitude 2*A* |cos [*π* (*ω*_1_ − *ω*_2_) *t*]| oscillates at a much slower frequency of |*ω*_1_ − *ω*_2_| (see Figure 3R). When considering multiple harmonic oscillators with their frequencies randomly sampled from a limited range, their superposition exhibits the characteristics of spindle oscillation. To confirm this, we extracted the oscillation frequencies of clusters in the TRN network at the MCP state (Figure S9), and used them as the frequencies of corresponding harmonic oscillators. Subsequently, we calculated the superposition of these harmonic oscillators. Our observations confirmed that the superposed activities indeed display the features of spindle oscillation, including rapid oscillation enveloped by waxing-and-waning amplitudes, as well as variations in spindle duration, amplitude, and inter-spindle interval (see Figure S9B and E). On the other hand, when we constructed harmonic oscillators using the oscillation frequencies of clusters in the phases of AFP and SFP, their superposed activities do not exhibit spindle oscillation, but rather demonstrate stochastic responses (Figure S9C and D) or regular oscillation (Figure S9A and F).

In the TRN network, we can regard a synchronized cluster as a harmonic oscillator. At the MCP state, all clusters represent distinct harmonic oscillators with slightly varying oscillation frequencies. Consequently, the collective activity of these clusters, reflected in the LFP, displays spindle oscillations, as described earlier (refer to Figure 2E and Figure 3D-F).

#### Spindle oscillations occur at the critical state

In the previous sections, we observed that spindle oscillations occur when the TRN network is in the MCP state, characterized by the formation of multiple synchronized clusters among the network’s neurons. Interestingly, this state aligns with the critical state of a dynamical system, a common feature of complex dynamical systems^74^. The critical state is known to emerge as a result of the interplay between synchronization and desynchronization forces within the system. When the synchronization force dominates, the system exhibits high order, with few synchronized clusters. Conversely, when the desynchronization force dominates, the system becomes highly disordered, lacking large-size clusters. The critical state arises when these two forces are balanced, resulting in the coexistence of multiple synchronized clusters within the network. Extensive theoretical studies have highlighted the significance of criticality as the optimal condition for large systems, including neuronal networks, to process information efficiently^74^.

In the TRN network, the synchronization force arises from electrical couplings between neurons, while the desynchronization force stems from neuronal heterogeneity. We have demonstrated that the interplay between these forces drives the TRN network into the MCP state, exhibiting the characteristic of criticality. In this section, we examine the correspondence between the MCP state and the critical state of the TRN network by computing relevant indicators of criticality.

Firstly, we examined the distribution of avalanche size in the TRN network. We began by calculating the spike times for all neurons and computing the average inter-spike interval (ISI) of these merged spike times. Using this ISI (typically 1-4 ms in our experiments) as the bin size, we divided the merged spike train into small bins and counted the number of spikes in each bin (see Figure 4A). We defined an avalanche as a series of consecutive bins, each containing at least one spike, separated by silent bins with no spike. The size of an avalanche was determined by the total number of spikes it contained (see the illustration in Figure 4A and STAR). It was observed that the distribution of avalanche sizes exhibited a power-law pattern exclusively in the MCP state (Figure 4D), suggesting criticality. In contrast, other states deviated from the power-law distribution (Figure 4C and E).

**Figure 4.**
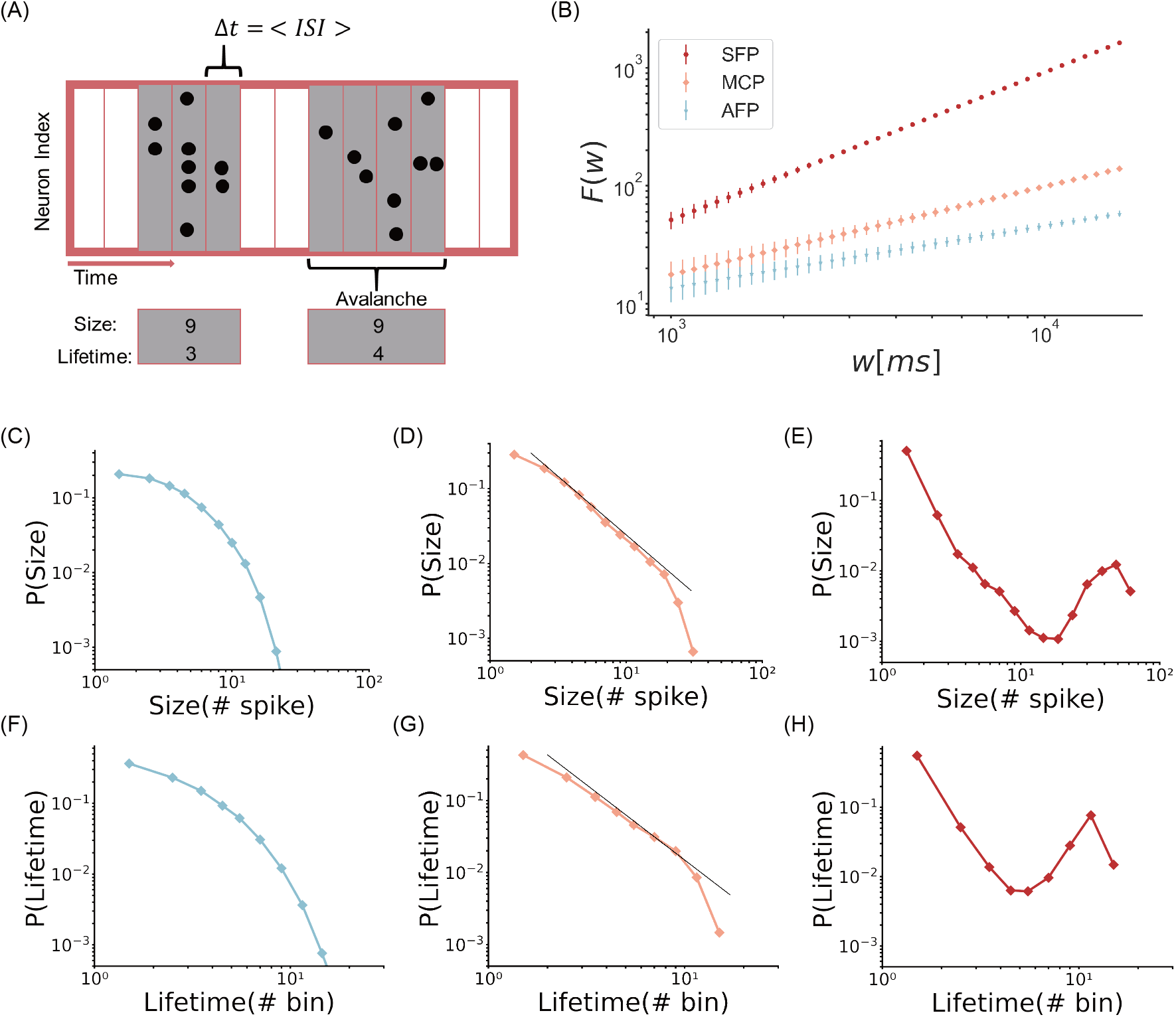
The criticality analysis of the spindle oscillation state. (A) The time is divided into small bins of equal length. The bin size Δ*t* is set to be the averaged inter-spike interval (ISI) of the merged spike train of all neurons, which is typically 1-4ms. An avalanche is defined as a sequence of bins with each of them containing at least one spike (marked by gray), and distinct avalanches are separated by silent bins. The size of an avalanche is the total number of spikes it contains, and the lifetime is the number of bins it spans.. (B) The DFA analysis of network oscillation behaviors. The x-axis represents the window width (*w*), while the y-axis the root-mean-squared fluctuation (*F* (*w*)) of the detrended time series of firing rate. The slope of the log-log plot is approximately 1.2, 0.85, and 0.5 in the state of SFP, MCP and AFP, respectively. Error bars represent the standard error of the mean (SEM) calculated from *n* = 10 independent simulations.. (C-E) The log-log plot of the distribution of avalanche size in three TRN states: the AFP state (C), the MCP state (D), and the SFP state (E).. (F-H) The log-log plot of the distribution of avalanche lifetime in three TRN states: the AFP state (F), the MCP state (G), and the SFP state (H).

Secondly, we measured the distribution of avalanche lifetime, which was defined as the number of bins an avalanche spans (see the illustration in Figure 4A and STAR). Only in the MCP state did the lifetime distribution exhibit a power-law pattern (Figure 4G), indicating criticality; while in other states, the power-law distribution was not satisfied (Figure 4F and H).

Thirdly, we assessed the long-range temporal correlation of oscillatory signals using detrended fluctuation analysis (DFA)^104^. DFA allows us to quantify the decay rate of auto-correlation in the network fluctuation *F* (*w*) (see STAR). If *F* (*w*) exhibits a slowly decaying auto-correlation structure, DFA yields a power-law form *F* (*w*) ∼ *w*^*α*^, where *α* ≈ 0.5 corresponds to a disordered pattern, 0.5 *< α <* 1 indicates criticality, and *α >* 1 reflects a highly correlated pattern. In the MCP state, we obtained 0.5 *< α <* 1, indicating that the network operates at the critical state (Figure 4B). Conversely, in AFP, *α <* 0.5, while in SFP, *α >* 1 (Figure 4B), suggesting deviations from criticality.

In summary, our results suggest that the TRN network operates at the critical state when spindle oscillations occur. However, it is important to note that the critical state alone is insufficient to generate spindle oscillation. For instance, if we counterbalance electrical coupling with stochastic noises instead of neuronal heterogeneity, we can achieve the criticality of the network state. Nevertheless, in such a case, all clusters within the network oscillate at the same frequency, and spindle oscillation will not manifest in the LFP. To generate spindle oscillation, the inclusion of neuronal heterogeneity is necessary since neuronal heterogeneity leads to clusters oscillating at slightly different frequencies, and their superposition gives rise to the waxing-and-waning shaped oscillation (refer to the analysis in Mathematical analysis of spindle generation).

## Conclusion and Discussion

The latest TRN experiments revealed that TRN neurons are highly heterogeneous and connected dominantly with electrical synapses. This suggests a reconsideration of the classical view of spindle generation via chemical synapses in the TRN network. In this work, by building a computational model capturing the updated properties of the TRN network, we re-examined the mechanism of spindle generation. We found that the waxing-and-waning spindle oscillations naturally emerge in the TRN network as a consequence of the competition between two driving forces: electrical synapses tending to synchronize neurons while heterogeneity tending to de-synchronize neurons. The interplay between two forces drives the TRN network to fall into a state where multiple synchronized clusters with slightly different oscillation frequencies coexist. The superposition of multiple local TRN clusters thus gives rise to spindle oscillation in the recorded LFP. Our research suggests that the TRN network operates at a critical state during spindle oscillations. This critical state represents a balance between order and disorder^74^ and allows the TRN network to integrate various inputs and coordinate neural activities across brain regions. These findings provide insights into spindle oscillation mechanisms and their potential role in learning and memory.

### Experimental evidence of the model

Our model has reproduced a number of experimental findings, including the statistical properties of the duration and the cycle number of spindle oscillations as observed in experiments^7,38,95^. It reconciles two conflicting experimental findings regarding the generation of TRN spindles. Steriade et al.^3,4,52^ found that the deafferented TRN *in vivo*, lacking thalamic and cortical projections, can generate and sustain spindle oscillations. On the other hand, von Krosigk et al.^31^ found that isolated TRN *in vitro*, devoid of any thalamic, cortical, and subcortical connections, fails to generate spindles. Our modeling supports both *in vivo* and *in vitro* experiments by highlighting that subcortical inhibitory projections *in vivo* serve as the excitatory drive for spindle generation, and this excitation arises due to the depolarized reversal potentials of TRN GABA_A_ receptors (*E*_GABA_ ≈ −45 mV found in^63^ and *E*_GABA_ = −62 ± 3.0 mV obtained in^64^).

Additionally, our model underscores the importance of electrical synapses and neuronal heterogeneity in spindle generation, corroborating recent experimental findings. For example, blocking electrical synapses suppresses spindle oscillation^105^, while reducing neuronal excitability by knocking out T-type Ca^2+^ channels diminishes spindle generation^38,79^.

Moreover, experimental studies have provided crucial insights into the synchronized behavior of TRN neurons under electrical couplings. Long et al.^66^ and Lee et al.^68^ observed clustered, synchronized responses in these neurons. Long et al.^66^ observed that when a subpopulation of TRN cells was activated by the nonspecific metabotropic glutamate receptor (mGluR) agonist ACPD, they generated synchronized rhythms in the 5-15 Hz range via electrical synapses, which is highly consistent with our modeling.

Finally, our model posits that neuronal heterogeneity is crucial for TRN spindle oscillation. Particularly, we incorporate two key aspects of this heterogeneity: variations in T-type Ca^2+^ conductance and the resulting diverse firing patterns, both of which are well-documented in experimental studies^81–83,102^. This heterogeneity is evident in the diverse morphology, neurochemistry, and electrophysiology of TRN neurons^78^. Research has identified distinct neuronal types^106^, synaptic projection patterns^107,108^, and neurochemical markers^37,109,110^, as well as genetic subtypes with varying firing properties^82,83,102^. These findings provide essential evidence supporting our model’s focus on neuronal heterogeneity.

### Predictions of the model

Our model produces predictions that can be tested through experiments (please also refer to STAR for the detailed analysis of model parameter influences on spindle oscillation properties). One of our key predictions is the significant role of electrical synapses in spindle generation and regulation. To validate this, we propose experiments that manipulate electrical synapse strength and observe the resulting effects on spindle oscillations. One such experiment involves measuring spindle oscillations in mice with Connexin 36 (Cx36) knockdown. Cx36 is the primary protein responsible for forming gap junctions in TRN [1,2]. Our model predicts that Cx36 knockdown mice will exhibit impaired spindle oscillations compared to wild-type mice.

Our model also suggests that neuronal heterogeneity, especially the variability in T-type calcium conductance and resultant firing patterns, is crucial for the desynchronization within the TRN network, leading to the characteristic waxing-and-waning spindle oscillations. To empirically test this hypothesis, we propose developing transgenic animal models with altered TRN neuronal heterogeneity and comparing their spindle oscillations to those of wild-type animals. A relevant study by Michael et al.^111^ reported diminished SK2 currents and neuronal bursting in TRN neurons of *Ptchd1*-knockout mice, with consequent reductions in sleep spindle counts. While this supports our model, the findings are limited by their *in vivo* context, which may include extraneous interactions between the thalamus and TRN. A more direct validation could be achieved by analyzing spindle oscillations in TRN slices from these knockout mice, thereby focusing solely on intrinsic TRN dynamics and eliminating external influences.

Additionally, our model postulates a universal mechanism for spindle-like oscillations, suggesting that the interplay between synchronizing and desynchronizing forces could generate similar oscillations across various neural circuits, not limited to the TRN. This hypothesis posits that waxing-and-waning oscillations could be a general feature of neural dynamics, observable across different brain regions and networks, reflecting a fundamental aspect of how neural systems transition between states of synchronization and desynchronization.

### Limitations of the study

Although our model successfully demonstrates the generation of spindle oscillations in an electrically coupled heterogeneous TRN network, there are several issues that require further investigation.

Firstly, the TRN is part of the larger corticothalamic system, influenced by excitatory inputs from thalamic and cortical areas and inhibitory inputs from subcortical regions. While our current focus was on the role of subcortical projections in spindle generation, the contribution of thalamo-cortical interactions is well-documented and crucial for the initiation, modulation, and termination of spindles^17^. Future enhancements of the model will incorporate these cortical influences to provide a more comprehensive understanding of spindle dynamics.

Secondly, the plasticity and asymmetry of electrical synapses between TRN neurons, which are believed to play significant roles in network oscillations, were not addressed in our current study^84,112,113^. In future work, we plan to extend our current model to investigate the roles of plasticity and asymmetry in spindle generation.

Thirdly, our analysis concentrates on thalamic spindles with frequencies in the slow range (7 − 12 Hz), yet spindle activity encompasses both slow and fast spindles, with the latter typically ranging from 12 − 15 Hz^114^. Our model primarily reproduces slow spindles, driven by the intrinsic oscillation frequencies of TRN neuron clusters. The rhythmic burst firing frequency of TRN neurons, influenced by voltage and temperature, generally matches this range. However, our simulated spindle frequencies are slightly slower than those recorded in experiments, where thalamic spindles often exceed 12 Hz^7^. This discrepancy may stem from our model’s exclusion of thalamic and cortical excitatory inputs, suggesting a need for further investigation into the factors that modulate spindle frequency in the thalamus. Future research should investigate the factors influencing thalamic spindle frequency to address this difference.

### Spindle oscillation at the critical state

Our modeling suggests that spindle oscillations emerge naturally in the electrically coupled TRN network when it self-organizes into the critical state due to the balance between electrical coupling and neuronal heterogeneity. Thus, we hypothesize that this critical state may play a functional role in efficient neural information processing within the thalamocortical system through two key aspects.

Firstly, the TRN has been suggested to play a strong functional control in attentional filtering^111,115–118^ and regulating oscillations^7,37,102,119^ in the brain. This functional role necessitates that the TRN network operates as a coherent whole while maintaining fine-grained control over individual neuronal units. Operating at criticality satisfies this requirement, as it allows the system to be exquisitely sensitive to inputs while enabling efficient signal propagation over long distances^74^. During spindle oscillations, this critical state likely allows for dynamic information processing from diverse subcortical and cortical inputs, with the coexistence of dynamically synchronized clusters acting as functional units to selectively gate or route information streams through the thalamus to the cortex.

Secondly, substantial experimental evidence from developmental and memory studies points to a fundamental role for sleep spindle oscillations in brain learning, plasticity, and memory^19–25^. Critical dynamics are known to enhance network adaptability and learning capabilities^120^. Therefore, the spindle critical state may optimize synaptic plasticity processes related to memory consolidation during sleep. By operating at criticality, the TRN network can efficiently coordinate and route relevant information streams from subcortical and cortical regions to support sleep-dependent memory consolidation and reorganization.

However, further experimental and theoretical studies are needed to directly test these hypothesized roles of the spindle critical state within the thalamocortical circuits (please refer to Section for limitations of the current model).

### The multifaceted origin of spindle generation

Recent experimental findings challenge the traditional perspective that spindle oscillations primarily originate from the thalamus^3,31,121,122^, suggesting that spindle oscillations may have diverse origins within the brain^33^. For example, evidence from EEG and MEG studies suggests a differentiation in their genesis: EEG spindles likely originate from matrix (nonspecific) thalamic nuclei, while MEG spindles stem from core (specific) thalamic pathways^123–125^. This distinction implies that various thalamic regions may contribute to specific types of spindle oscillations. Furthermore, recent pharmacological studies in humans have shown that ion channel antagonists differentially affect fast and slow spindles^114^. Notably, the administration of a Ca^2+^ blocker selectively suppresses fast spindles, while leaving slow spindles intact. These findings collectively reinforce the hypothesis that spindle oscillations arise from multiple, distinct neurophysiological mechanisms.

Although the model is based on the TRN site, our modeling suggests a general mechanism for the genesis of spindle-like oscillations, where self-organized synchronization and desynchronization forces create the critical state. In this state, neuronal clusters dynamically appear and disappear, corresponding to the waxing-and-waning phase of spindles, respectively. This mechanism may extend beyond the TRN, potentially implemented through various neural circuits. For instance, the interplay between cortical and thalamic inputs also plays a significant role in this process. For example, cortical feedback and thalamic projections modulate TRN neurons during the cortical slow oscillation cycles, where the upstate phases enhance synchronization and the downstate phases promote desynchronization^40,126^.

### Related works

Various mechanisms for spindle oscillations have been proposed previously. As noted in introduction, much of the previous research has focused on the role of mutual inhibition via chemical synapses between TRN neurons in generating spindles^34,59–61^. Another significant line of computational work explores the “ping-pong” interaction between TC and TRN neurons. These studies highlight how different connectivity patterns^35,124,127^, ion channel dynamics^36^, neuromodulation effects^128^, and short-term plasticity dynamics^129^ can lead to different types of spindle oscillations. Moreover, direct spindle-like modulation inputs have been investigated as another mechanism influencing spindle activities. For example, studies by Yazdanbakhsh et al.^130^ have demonstrated how spindle-like inputs can shape the activity patterns in the TRN. The most relevant work to our study is a pioneering model developed by Fuentealba et al.^105^, in which TRN neurons were interconnected via both inhibitory chemical synapses and electrical synapses. However, in Fuentealba et al.^105^, spindles arise from the mutual inhibition and rebound bursting of TRN neurons.

## Supplemental information index

Figures S1-S12 and their legends in a PDF

## Acknowledgments

This work was supported by the Science and Technology Innovation 2030-Brain Science and Brain-inspired Intelligence Project (No. 2021ZD0200204, SW), the Young Scientists Fund of the National Natural Science Foundation of China (No. 32400937, CMW), the China Postdoctoral Science Foundation (No. 2024M750076, CMW), and the Postdoctoral Fellowship Program of CPSF (No. GZC20230106, CMW). The authors thank all members of the lab for their support.

## Author contributions

Conceptualization, S.L., C.W., and S.W.; Methodology, S.L., C.W., and S.W.; Writing, S.L., C.W., and S.W. Funding Acquisition, S.W.; Supervision, S.W.

All Cell Press **life science** journals and **iScience** use the **STAR Methods** format for reporting materials and methods. Other Cell Press journals use an **experimental procedures** format; for those journals, please **delete** the contents of this page from the template and refer instead to the **experimental procedures** template on page 2.

The following headings are mandatory and must be included exactly as they are written here: Key resources table, Resource availability (including subheadings Lead contact, Materials availability, and Data and code availability), and Method details. Please do not alter the mandatory headings. All other headings are optional.

Please refer to the full STAR Methods guide for authors for further information:

- *Cell* authors should download the guide for *Cell* authors available here
- All other journal authors should use this version of the guide

## STAR METHODS

### Key resources table

*To create the KRT, please use the KRT webform or the Word template and upload this file separately*.

### Resource availability

#### Lead contact

Further information and requests for resources and reagents should be directed to and will be fulfilled by the lead contact, Si Wu.

#### Materials availability

The study did not generate new unique reagents.

### Data and code availability

- All data reported in this paper can be accessed through the open-access repository at DOI 10.5281/zenodo.13377848.
- The source code, processed data, and statistical analyses supporting the findings of this study are available at the open-access repository https://github.com/shangyangli/TRN-spindl and have been archived at DOI 10.5281/zenodo.13377848.
- Any additional information required to reanalyze the data reported in this paper is available from the lead contact upon request.

### Experimental model and study participant details

This study does not involve any experimental models or human participants. Instead, a computational neuroscience model was developed and used to conduct the research. Therefore, there are no species, genotypes, ages, sexes, or maintenance and care information to report.

## Method details

### Network structure

To mimic a 1 × 1 mm of TRN slice, we modeled a tier of TRN network which contains 100 (10 × 10) neurons (see Figure 1A). The TRN network is placed in a two-dimensional grid with a periodic boundary, in which neurons are electrically coupled with each other by a Gaussian probability connection pattern. The connection probability for each pair of neurons (*i, j*) obeys

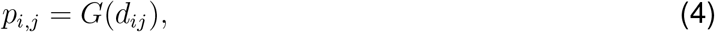

where *d*_*ij*_ is the Euclidean distance between two neurons, and *G* is a 2-dimensional Gaussian function centered at 0 with the standard deviation *σ*, and *σ* controls the connection width of each neuron.

In addition to the electrical couplings between neurons, each neuron may receive two other kinds of synaptic projections (see Figure 1A). To mimic the physiological condition of the TRN network in *vivo*, a neuron receives the inhibitory projections from the subcortical structures^50,51,118^. This inhibitory projection is modeled as a train of Poisson spikes conveying currents onto the neuron with GABA_A_ synapse. To model the synaptic connections from the thalamus and cortex in the awake state, a neuron receives a spike train with excitatory AMPA synapse. Particularly, the firing rates of both excitatory and inhibitory background inputs used in our model are 50 Hz.

### TRN neuron model

For the convenience of analysis, we used the previously developed reduced TRN neuron model^75^. It is biologically grounded, capable of reproducing characteristic TRN bursts and tonic discharges, and allows easy dynamical analysis.

Specifically, the TRN neuron is modeled as a single-compartment Hodgkin-Huxley model, with a minimal set of active channels that are sufficient to produce the tonic and bursting spiking dynamics. It contains three active ionic currents: a fast sodium current (*I*_*Na*_) and a delayed rectifier (*I*_*K*_) for spiking generation and a low-threshold Ca^2+^ current (*I*_*T*_) for rebound bursting. Moreover, it has two passive leaky channels: a potassium leaky current (*I*_*KL*_) for modeling the neuromodulation effect, and a leaky current (*I*_*L*_).

The dynamics of the TRN neuron model is given by

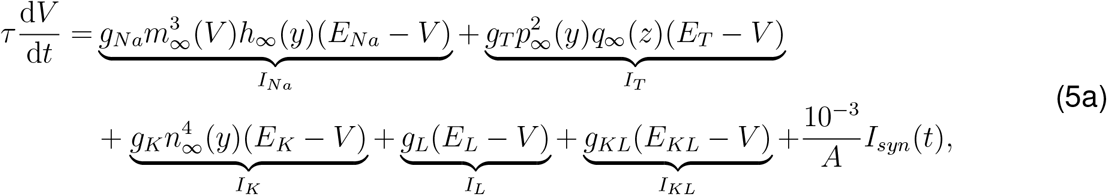

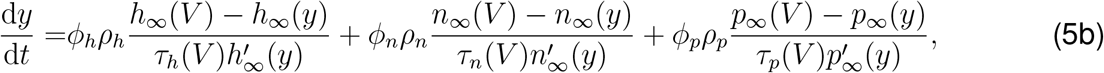

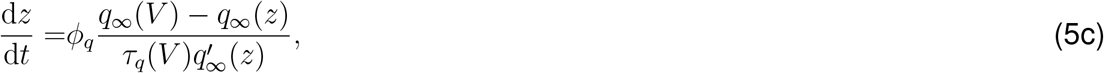

where *τ* = *C/ρ*_*V*_ is the membrane time constant, *I*_*syn*_ is the received synaptic current that normalized by the membrane area *A* = 1.43 × 10^*−*4^ cm^2^, *I*_*Na*_, *I*_*K*_ and *I*_*T*_ are the currents mediated by the sodium, potassium and low-threshold calcium channels, respectively, *I*_*L*_ is the leakage current, *I*_*KL*_ is the potassium leaky current controlled by neuromodulators like acetylcholine and norepinephrine, *C* = 1 *µ*F*/*cm^2^ is the membrane capacitance, *ρ*_*V*_ is the reduction coefficient of the membrane potential *V* ^75^, *y* and *z* are two reduced equivalent potentials^75^, *ρ*_*h*_, *ρ*_*n*_ and *ρ*_*p*_ are reduction coefficients^75^, *ϕ*_*h*_ = 1, *ϕ*_*n*_ = 1, *ϕ*_*p*_ = 6.9 and *ϕ*_*q*_ = 3.7 are temperature factors, *τ*_*x*_(*x* ∈ {*h, n, p, q*}) are time constant functions of each variable, and *x*_∞_(*x* ∈ {*h, n, p, q*}) are steadystate functions. The detail expressions of reduction coefficients (including *ρ*_*V*_, *ρ*_*h*_, *ρ*_*n*_, *ρ*_*p*_, *ρ*_*q*_), time constant functions (including *τ*_*h*_, *τ*_*n*_, *τ*_*p*_, *τ*_*q*_), and steady-state functions (including *h*_∞_, *n*_∞_, *p*_∞_, *q*_∞_) can be obtained in^75^.

For the sodium current *I*_*Na*_, we set the maximal conductance *g*_*Na*_ = 100 mS*/*cm^2^, the reversal potential *E*_*Na*_ = 50 mV, and the spike adjusting threshold 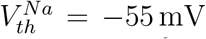 mV. For the potassium current *I*_*K*_, we set 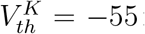 mV, *E*_*K*_ = −100 mV, and *g*_*K*_ = 10 mS*/*cm^2^. For the low-threshold calcium current *I*_*T*_, we set *g*_*T*_ = 2.25 mS*/*cm^2^, and 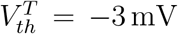 mV. For the leakage current *I*_*L*_, we used the leakage channel conductance *g*_*L*_ = 0.06 mS*/*cm^2^ and the leakage reversal potential *E*_*L*_ = −70 mV. For the potassium leaky current *I*_*KL*_, we set the potassium leakage channel conductance *g*_*KL*_ = 0.0065 mS*/*cm^2^ and the reversal potential *E*_*KL*_ = −100 mV.

The synaptic current of the TRN neuron *i* is decomposed into three components:

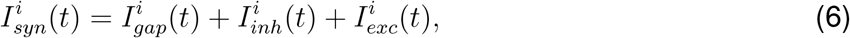

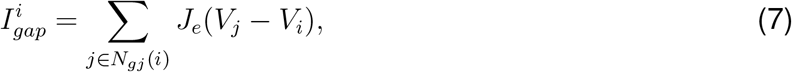

where *I*_*gap*_ is the electrical synapse current, *I*_*inh*_ the inhibitory GABA_*A*_ current, and *I*_*exc*_ the excitatory AMPA current. They are described in the following.

### Electrical synapse model

The electrical synapse current flowing into the TRN neuron *i* is computed as
where *J*_*e*_ is the conductance of electrical synapses, and *N*_*gj*_(*i*) denotes the collection of neurons that are electrically coupled with the neuron *i*.

### Subcortical inhibitory synapse model

The inhibitory synaptic current 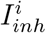 to the neuron *i* from the transmission of a subcortical spike is computed as the GABA_*A*_ synapse model. Specifically, it is modeled as:

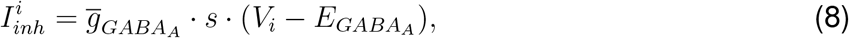

where 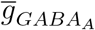 denotes the maximum GABA_*A*_ synaptic conductance, 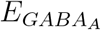 the synaptic re-versal potential, and *s* the fraction of open channels. Moreover, *s* is modeled by the first-order activation scheme

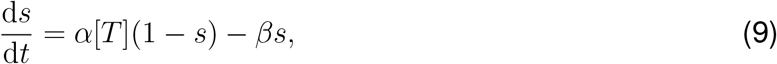

where [*T* ] is the concentration of neurotransmitter modeled as a brief pulse (0.3 ms duration, 0.5 mM amplitude) after pre-synaptic neuron generating an action potential^36^, and *α* and *β* are the channel forward and backward binding rates given by *α* = 10 ms^*−*1^ and *β* = 0.18 ms^*−*1^, respectively^36,129,131^.

### Thalamic and cortical excitatory synapse model

The excitatory synaptic current 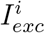 to the neuron *i* when receiving a thalamic or cortical spike is computed as the AMPA synapse model. Specifically, the AMAP current is modeled as

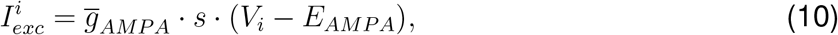

where 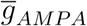 denotes the maximum AMPA synaptic conductance, and *E*_*AMPA*_ the synaptic reversal potential. Similar to GABA_*A*_ model, *s* is modeled by Equation 9, except that *α* = 10.5 ms^*−*1^ and *β* = 0.166 ms^*−*1^ ^36,129,131^.

### Simulation tool

The network model was coded with the BrainPy library in Python^132,133^. All simulations were performed by using the fourth-order Runge-Kutta method (RK4) with a fixed time step of 0.01 ms. A shorter simulation time step does not change the results. The model source code is available on the GitHub.

### Local field potential

We measured the local field potential by averaging the membrane potentials across the whole TRN neuron group^76^, which is written as,

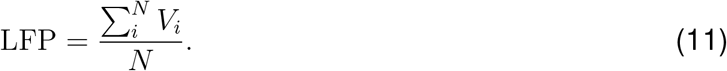

We then used a finite impulse response (FIR) filter (mne.filter.filter_data(), a built-in function implemented in MNE-Python toolbox^134^) to calculate the broadband filtered LFP signal (0.5 ∼30 Hz) LFP_bf_ and the spindle band filtered signal (6 ∼10 Hz) LFP_spin_. The power spectrum of the LFP was obtained by a fast Fourier transform (FFT) on LFP_bf_. The peak spectral frequency was chosen as the network oscillation frequency.

In addition to the standard method of calculating the local field potential (LFP) by averaging the membrane potentials across the whole TRN neuron group, we also employed a spatial lowpass filter to compute the LFP. This method takes into account the spatial distribution of neurons and their respective distances, providing a more nuanced representation of the neuronal activity^135,136^.

The spatial lowpass filter method involves the following steps:

#### 1 Distance Matrix Calculation

We first calculate the spatial distance matrix, *D*, where each element *D*_*ij*_ represents the Euclidean distance between neuron *i* and neuron *j*. The positions of neurons are assumed to be in a two-dimensional space.

#### 2 Weight Matrix Calculation

A Gaussian function is used to calculate the weight matrix, *W*, based on the distances. The weight for each pair of neurons *i* and *j* is given by:

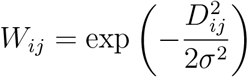

where *σ* is a parameter that controls the spread of the Gaussian function. The weights are then normalized so that the sum of the weights for each neuron is equal to 1:

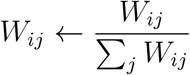

#### 3 LFP Calculation

Using the weight matrix, the LFP is calculated as a weighted sum of the membrane potentials, *V* :

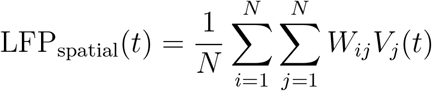

where *N* is the total number of neurons, and *V*_*j*_(*t*) is the membrane potential of neuron *j* at time *t*.

The comparison between the standard LFP method and the spatial lowpass filter method is illustrated in Supplementary Materials (Figure S12).

### Automatic spindle detection

Spindles are automatically detected, with the method inspired by the A7 algorithm described in^137^ and the YASA spindle algorithm presented in^138^. Specifically, four thresholds are used:

#### Relative spindle power RelSpinPow

The first threshold is computed as the relative power in the spindle frequency range, which aims to ensure the increase of the power is specific to the spindle oscillation range and not due to a global oscillation power increase. The threshold is set to RelSpinPow ≥ 0.9.

#### Root mean square RMS

In order to detect the energy power increase in the spindle frequency band, the moving RMS of LFP_spin_ with a window size of 100 ms and a step size of 50 ms is used. The threshold is exceeded whenever a sample has a RMS ≥ RMS_threshold_, where RMS_threshold_ = RMS_mean_ + 0.1RMS_std_.

#### Spindle correlation SpinCorr

In order to characterize the causal influence of LFP_spin_ power increase on LFP_bf_ rise, SpinCorr is computed by the Pearson correlation coefficient between LFP_spin_ and LFP_bf_ with a moving sliding window of 300 ms and a step of 100 ms. The threshold is exceeded whenever a sample has a correlation value SpinCorr ≥ 0.9.

#### Gaussian kernel fitting errors err_*p*_

When the above three parameters exceed their respective thresholds, a candidate spindle event can be selected. However, the morphologies of such events extracted by three thresholds may not be waxing and waning. Therefore, we fit the cycle peaks in each spindle candidate with a Gaussian kernel 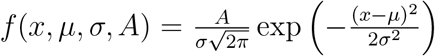 to filter out the false alarm event. The standard deviation errors (err_*p*_) on the parameters {*A, µ, σ*} after curve fitting are used for the threshold judgment. Once one of the errors exceeds 10, we reject this candidate event.

### Cross correlation

The cross-correlation between a pair of neurons is calculated as follows. Denoting the spike trains of a pair of neurons to be *S*_*i*_(*t*) and *S*_*j*_(*t*), for *S*_*i*_(*t*), *S*_*j*_(*t*) = 0 or 1, and *t* = 1, 2, …, *T*, the CC between the neuron pair is calculated to be

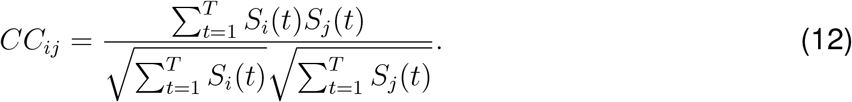

For a network, the overall cross-correlation is given by,

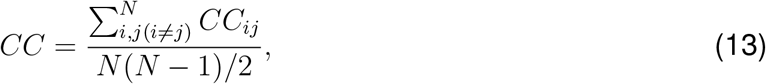

where the summation runs over all neuron pairs.

### Coupling coefficient

To assess the strength of the electrical synapse connection, we utilized the steady-state coupling coefficient (CC) as a measure. This involved injecting long polarizing current pulses into one of the paired neurons. The injection of current induced a voltage drop in the presynaptic cell (V_pre_), which typically caused a change in the membrane potential of the coupled postsynaptic cell (V_post_). The duration of the current pulses was intentionally extended to overcome the initial attenuation of the membrane potential caused by the passive properties of the postsynaptic membrane’s filtering characteristics. Consequently, we were able to measure voltage changes in a stable state. Quantitatively, the strength of coupling was determined by calculating the coupling coefficient, defined as the ratio between the voltage deflections observed in the post- and presynaptic cells:

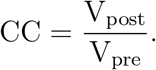

### Critical state analysis

#### Data Preprocessing for Avalanche Analysis

Following the classical analysis of criticality in spiking activity^139^, we counted the number of spikes in each bin of length Δ*t* (measured in milliseconds) for the merged spike train of all neurons. An avalanche is defined as a sequence of consecutive bins containing spikes, separated from other avalanches by silent bins. In this study, we defined the size of an avalanche *S* as the total number of spikes it contains, and the lifetime *T* as the number of bins it spans. We selected the average inter-spike interval of the merged spiking train as the bin size Δ*t*.

#### Critical exponents estimation

First, we inspected the distributions of avalanche size and lifetime. For distributions exhibiting characteristics of a power law, we used a doubly truncated algorithm based on maximum likelihood estimation^140,141^ for analysis. Denote *x* the size or life-time of an avalanche. We fit a discrete power-law distribution:

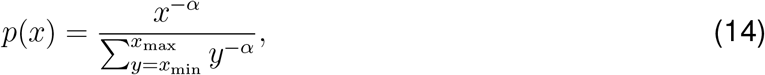

where *x*_min_ is the smallest avalanche size or lifetime, and *x*_max_ is the upper bound. Assume identical and independent distribution of avalanches, the value of *α* can be calculated by maximizing the likelihood function, which is,

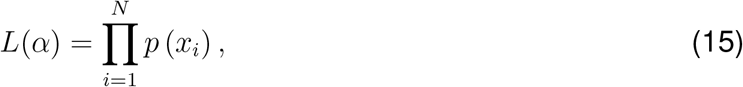

where *N* is the number of avalanches.

To assess the goodness of fit and determine if a power law adequately describes the data, we used several evaluation techniques^142^. First, we employed bootstrapping and the Kolmogorov-Smirnov test to generate a *p*-value for an individual fit. Second, we used the log-likelihood ratio to compare two fits and identify the better one. The *p*-value from the Kolmogorov-Smirnov test represents the significance level of the fit. Comparing the log-likelihood ratio between two candidate distributions helps determine if the data is more likely to follow one distribution over the other. A positive log-likelihood ratio indicates that the data is more likely to adhere to the first distribution, while a negative value suggests a greater likelihood for the second distribution^142^.

#### Detrended Fluctuation Analysis

In analyzing the scale-invariant long-range time correlation, we employed detrended fluctuation analysis (DFA)^104^. This method works as follows: 1) we first calculated an integrated zero-mean version of the original time series *z*, denoted as *y*(*k*), by summing the deviations from the mean up to the *k*-th point, i.e., 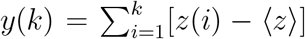; 2) we then divided the entire time period into consecutive non-overlapping window of width *w* (in milliseconds or seconds, as appropriate) and fit a local linear trend *y*_*w*_ in each window; 3) finally, we calculated the fluctuation *F* (*w*) at a given time-scale *w* using the following equation:

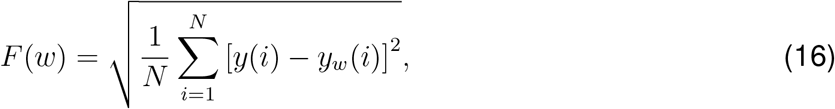

where *N* is the total number of time windows.

The scaling exponent *α* is determined by linearly fitting log *F* (*w*) vs log *w*. The interpretation of *α* is as follows:

- *α <* 0.5: anti-correlation in the time-series.
- *α* = 0.5: absence of correlation (white noise).
- 0.5 *< α <* 1: positive correlation in the time series, indicating that the neural network operates in the critical regime, where small fluctuations can propagate and lead to complex, system-wide response.
- *α* ≈ 1: 1/f-noise, pink noise.
- *α >* 1: positive correlation often seen in highly structured systems.

### Analysis of the TRN neuron model

Diverse firing patterns, including different degrees of bursting^143–146^ and tonic spiking^80,119,143,147^, have been observed in TRN neurons *in vivo* and *in vitro*. To understand the neuronal mechanism of TRN’s characteristic burst and tonic discharges, we previously developed a reduced three-variable TRN neuron model^75^ which allows us to perform bifurcation analysis of its spiking dynamics (see the following text). Despite its simple form, this neuron model can reproduce the discharge patterns as seen in TRN oscillatory activities^4,7,31,38,52,145,148^ (see Figure S1).

Our reduced model shows different degrees of rhythmic bursts under different resting potentials (Figure S1A). Under the depolarized states (like *V*_*r*_ = −61 or −65 mV), the cell exhibits burst firings with the spindle frequency (7-15 Hz). While hyperpolarization of neurons to −70 and −75 mV results in a decrease in interburst frequency and a substantial increase of spike number in a burst. Similarly, this neuron model shows consistent tonic spiking (Figure S1B) as seen in biological observations^7^. More hyperpolarization of the neuron needs a larger size of excitatory current to elicit a TRN spike. What’s more, as shown in Figure S1C, our single neuron model can exhibit different degrees of rebound bursting as observed in biological experiments^82,83,146^. Specifically, neurons with larger *g*_*T*_ conductance show higher intrinsic bursting, while small *g*_*T*_ conductance results in a few spikes per rebound burst.

Diverse discharge patterns of the TRN neuron occur conditionally, depending on the conductance of low-threshold Ca^2+^ current *g*_*T*_, the conductance of potassium leaky current *g*_KL_, and the external inputs. The intrinsic bursting oscillations in the TRN neuron are more obvious when *g*_*T*_ becomes larger (Figure S1D). It can oscillate with small values of *g*_KL_, i.e., at high resting membrane potential, when *g*_*T*_ is large, implying that the larger size of *g*_*T*_ increases the intrinsic excitability of the model. However, when the cell has small values of *g*_*T*_, the intrinsic oscillation of the model disappears no matter how *g*_KL_ changes (Figure S1E). Instead, the subthreshold oscillation as reported in the experiments^66^ occurs when *g*_KL_ lies in the intermediate value regime.

### Superposition of harmonic waves

We superpose two simple harmonic waves with the same direction and frequency. Suppose the two simple harmonic waves with the same direction and frequency are:

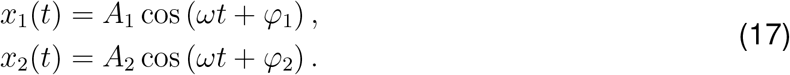

Their summation is calculated to be,

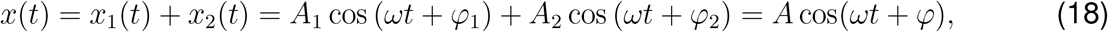

where,

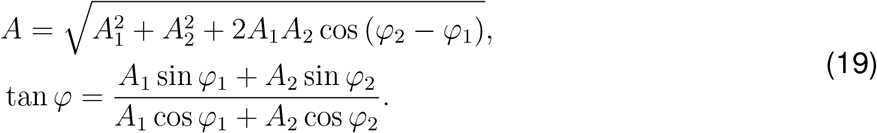

It can be seen that the superposition of two simple harmonic waves with the same direction and frequency is still simple harmonic.

We next examined the superposition of two simple harmonic waves with slightly different frequencies. Consider the two simple harmonics are:

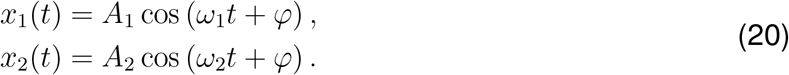

Further assume *A*_1_ = *A*_2_ and |*ω*_1_ − *ω*_2_| *<< ω*_1_ + *ω*_2_, the superposition of two waves are:

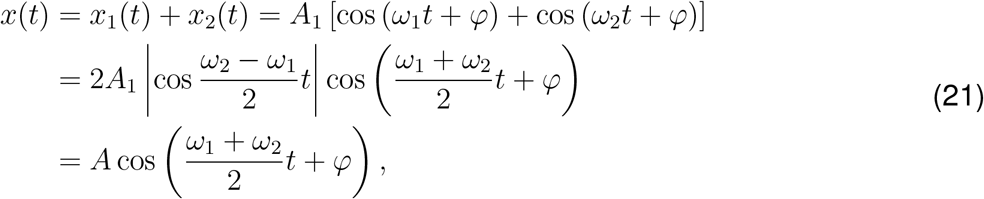

where *A* = 2*A*_1_| cos(*ω*2 − *ω*1)t*/*2| is the amplitude of the superimposed wave. Since |*ω*_2_ − *ω*_1_ ≪ *ω*_1_ + *ω*_2_, the superposed signal *x*(*t*) exhibits the wax-and-waning shape.

### The influence of model parameters on spindle oscillation properties

To understand the properties of spindles generated in the TRN network, we quantitatively studied the influence of network parameters on the simulated spindle activities. We focused on the electrical synapse strength (corresponds to the average CC, see Figure S2A, D, G and Figure S3 A1-A6), the neuromodulation control level (corresponds to the potassium leaky conductance *g*_KL_, see Figure S2B, E, H and Figure S3B1-B6), and the heterogeneity of neuronal firing patterns (corresponds to the variance of Ca^2+^ channel conductance *g*_*T*_, see Figure S2C, F, I and Figure S3C1-C6).

We analyzed various metrics of spindle oscillation using the YASA spindles detection algorithm^138^. These metrics provide valuable insights into the characteristics of spindles. The specific metrics we examined include:

- Frequency: This metric represents the median instantaneous frequency of the spindle, measured in Hz. It is derived from a Hilbert transform of the filtered signal.
- Duration: This metric indicates the duration of spindle oscillation, measured in seconds.
- Amplitude: The amplitude metric represents the peak-to-peak amplitude of the detrended spindle in the raw data. It is measured in microvolts (*µ*V).
- AbsPower: This metric quantifies the median absolute power of the spindle, measured in log 10 *µV* ^2^. It is calculated from the Hilbert-transformed filtered signal.
- Oscillations: This metric denotes the number of oscillations, which is equivalent to the number of positive peaks in the spindle.
- Peak: This metric indicates the time at the most prominent spindle peak, measured in seconds.
- RelPower: Similar to the AbsPower metric, this metric represents the median absolute power of the spindle in log 10 *µV* ^2^. It is derived from the Hilbert-transformed filtered signal.
- RMS: This metric stands for root-mean-square and is measured in microvolts (*µ*V).
- Symmetry: This metric indicates the location of the most prominent peak of the spindle, normalized on a scale from 0 (start) to 1 (end). An ideal value for this metric is close to 0.5, indicating that the most prominent peak is located halfway through the spindle.

The average CC of the TRN network was observed to have a great impact on the spindle oscillation frequency (Figure S2A) and amplitude (Figure S2G). With the increasing average CC, the TRN generates spindles at a lower frequency (Figure S2A). Higher CC values induce more synchronized network spiking, resulting in spindle oscillations with a higher amplitude (Figure S2G) and stronger absolute power in simulated LFPs (Figure S3A). The increased oscillation amplitude and power are associated with higher RMS values of the spindle signal (Figure S3A5), indicating that larger CC results in greater variance in spindle LFP activity. Moreover, increasing CC slightly increased the duration (Figure S2B) and the number of oscillation cycles (Figure S3A2).

The effect of neuromodulation on spindle rhythms showed different profiles. Increasing the neuromodulation level, i.e., decreasing *g*_KL_ and increasing the resting membrane potential of each TRN neuron, augmented the oscillation frequency of spindle activity (Figure S2B), as depolarized resting potentials increased single-neuron burst frequencies (see Figure 1). Meanwhile, the depolarized membrane potential inactivated the Ca^2+^ channel and reduced burst strength in individual TRN neurons, leading to a reduction in oscillation amplitude (Figure S2H), oscillation power (Figure S2B1), and LFP signal variance (Figure S3E) with smaller values of *g*_KL_. Unlike the network average CC, no consistent impact of *g*_KL_ on spindle duration (Figure S2B5), oscillation cycles (Figure S3B2), and relative oscillation powers (Figure S3B4) was observed.

Increasing the variance of *g*_*T*_ in TRN neurons had the opposite effect on spindle activity compared to increasing CC. A higher *g*_*T*_ variance means more TRN neurons exhibit higher *g*_*T*_ values, leading to more frequent bursting (Figure 1). As a result, increasing *g*_*T*_ variance led to spindles with higher oscillation frequencies (Figure S2C). However, the increase in *g*_*T*_ variance also led more neurons to have lower *g*_*T*_ values, which caused these neurons to spontaneously oscillate at subthreshold levels. Consequently, the TRN network generated spindle activities with lower amplitude as *g*_*T*_ variance increased (Figure S2I). Correspondingly, we observed that absolute oscillation power and RMS decreased with higher *g*_*T*_ variance (Figure S3C1 and C5). In contrast to electrical coupling strength, increased heterogeneity among TRN neurons slightly decreased spindle oscillation duration (Figure S2F) and oscillation cycles (Figure S3C2). Interestingly, a substantial increase in relative oscillation power at the spindle band was observed (Figure S3C4), implying that spindle activity dominated network oscillations with increased TRN cell heterogeneity.

Overall, these three parameters were observed to modulate different aspects of TRN spindle rhythms. However, no consistent influence of these parameters on spindle oscillation shape, including oscillation peak and symmetry, was observed (see Figure S3A3, B3, C3 and Figure S3A5, B5, C5). These properties may be shaped by other network parameters and warrant further investigation.

